# Near-optimal combination of disparity across a log-polar scaled visual field

**DOI:** 10.1101/589937

**Authors:** Guido Maiello, Manuela Chessa, Peter J. Bex, Fabio Solari

**Affiliations:** Department of Experimental Psychology, Justus-Liebig University Gießen, Gießen, Hesse, Germany; Department of Psychology, Northeastern University, Boston, Massachusetts, USA; Department of Informatics, Bioengineering, Robotics and Systems Engineering, University of Genoa, Genoa, Italy

## Abstract

The human visual system is foveated: we can see fine spatial details in central vision, whereas resolution is poor in our peripheral visual field, and this loss of resolution follows an approximately logarithmic decrease. Additionally, our brain organizes visual input in polar coordinates. Therefore, the image projection occurring between retina and primary visual cortex can be mathematically described by the log-polar transform. Here, we test and model how this space-variant visual processing affects how we process binocular disparity, a key component of human depth perception. We observe that the fovea preferentially processes disparities at fine spatial scales, whereas the visual periphery is tuned for coarse spatial scales, in line with the naturally occurring distributions of depths and disparities in the real-world. We further show that the visual field integrates disparity information across the visual field, in a near-optimal fashion. We develop a foveated, log-polar model that mimics the processing of depth information in primary visual cortex and that can process disparity directly in the cortical domain representation. This model takes real images as input and recreates the observed topography of disparity sensitivity in man. Our findings support the notion that our foveated, binocular visual system has been moulded by the statistics of our visual environment.

**Author summary:** We investigate how humans perceive depth from binocular disparity at different spatial scales and across different regions of the visual field. We show that small changes in disparity-defined depth are detected best in central vision, whereas peripheral vision best captures the coarser structure of the environment. We also demonstrate that depth information extracted from different regions of the visual field is combined into a unified depth percept. We then construct an image-computable model of disparity processing that takes into account how our brain organizes the visual input at our retinae. The model operates directly in cortical image space, and neatly accounts for human depth perception across the visual field.

## Introduction

Humans employ binocular disparities, the differences between the views of the world seen by our two eyes, to determine the depth structure of the environment. Additional complexity in our estimate of the depth structure arises because spatial resolution is not uniform across the visual field. Instead, our visual system is space-variant: the foveae of both our eyes are sensitive to fine spatial detail, while vision in our periphery is increasingly coarse. Therefore, when humans look at an object, the eyes are rotated so that the high-resolution foveae of both eyes are pointed at the same location on the surface of the object. The fixated object will extend into our binocular visual field by a distance proportional to the object’s size, and over this area we will experience small stereoscopic depth changes, arising from retinal disparities due to the surface structure and slant or tilt of the fixated object. The world beyond the fixated object in our peripheral visual field will typically contain objects at a range of different depths. Consequently we will experience a greater magnitude and range of binocular crossed (closer than the fixated horopter) and uncrossed (farther than the fixated horopter) disparities [1]. It has been proposed that the visual system may process disparity at different spatial scales along separate channels [2], analogous to the channels selective for luminance differences at different spatial frequencies [3]. Using a variety of paradigms, several authors have provided evidence for at least two [4-10] or more [11-13] spatial channels for disparity processing.

Given that our visual world contains small, fine disparities near the fovea and larger coarse disparities in our peripheral visual field, we might analogously expect sensitivity to disparity to vary across the visual field. Based on differences in experience during development, different regions of our visual field might therefore be expected to be optimized to process disparity at different spatial scales [14]. We test this hypothesis by measuring disparity sensitivity to annular stimuli spanning rings of different retinal eccentricity across the visual field in human participants. We hypothesize that as eccentricity increases from fovea to periphery, the tuning of depth sensitivity should shift from fine to coarse spatial scales. We also hypothesize that peak sensitivity to stereoscopic disparity should also decrease as eccentricity increases, following the general decrease in visual sensitivity observed in the visual periphery [15].

If indeed different visual field eccentricities preferentially process disparities at different spatial scales, then how does the visual system combine depth information processed throughout our visual field to recover the depth structure of the environment? If disparity information is integrated across different regions of the visual field, then sensitivity for full field stimuli should be better than for stimuli spanning smaller areas of the visual field. We test whether this integration process is optimal according to a maximum-likelihood estimation (MLE) principle [16-22].

Next, we construct a model. Prince and Rogers [14] were the first to suggest that disparity sensitivity across the visual field may be related to M-scaling. Gibaldi et al [23] even suggest that the specific pattern of cortical magnification might be a consequence of how we visually explore the naturally occurring distribution of real-world depths. Therefore, we implement a simple, neurally-inspired model of disparity processing, in which we include a critical log-processing stage that mimics the transformation between retinal and cortical image space [24-26]. A unique advantage of this approach is that disparity can be computed and analyzed directly in the cortical domain [27]. We have previously shown that this approach can account for motion processing throughout the visual field of human participants [28]. Here, we examine whether log-polar processing can also account for human disparity processing across the visual field.

## Results

Figure 1a shows the stimuli we employed to psychophysically assessed disparity sensitivity in the central (red, 0-3 deg), mid peripheral (green, 3-9 deg), far peripheral (blue, 9-21 deg), and full (black, 0-21 deg) visual field of human observers (see detailed descriptions of stimuli and experimental procedures in the Methods section).

**Fig 1.**
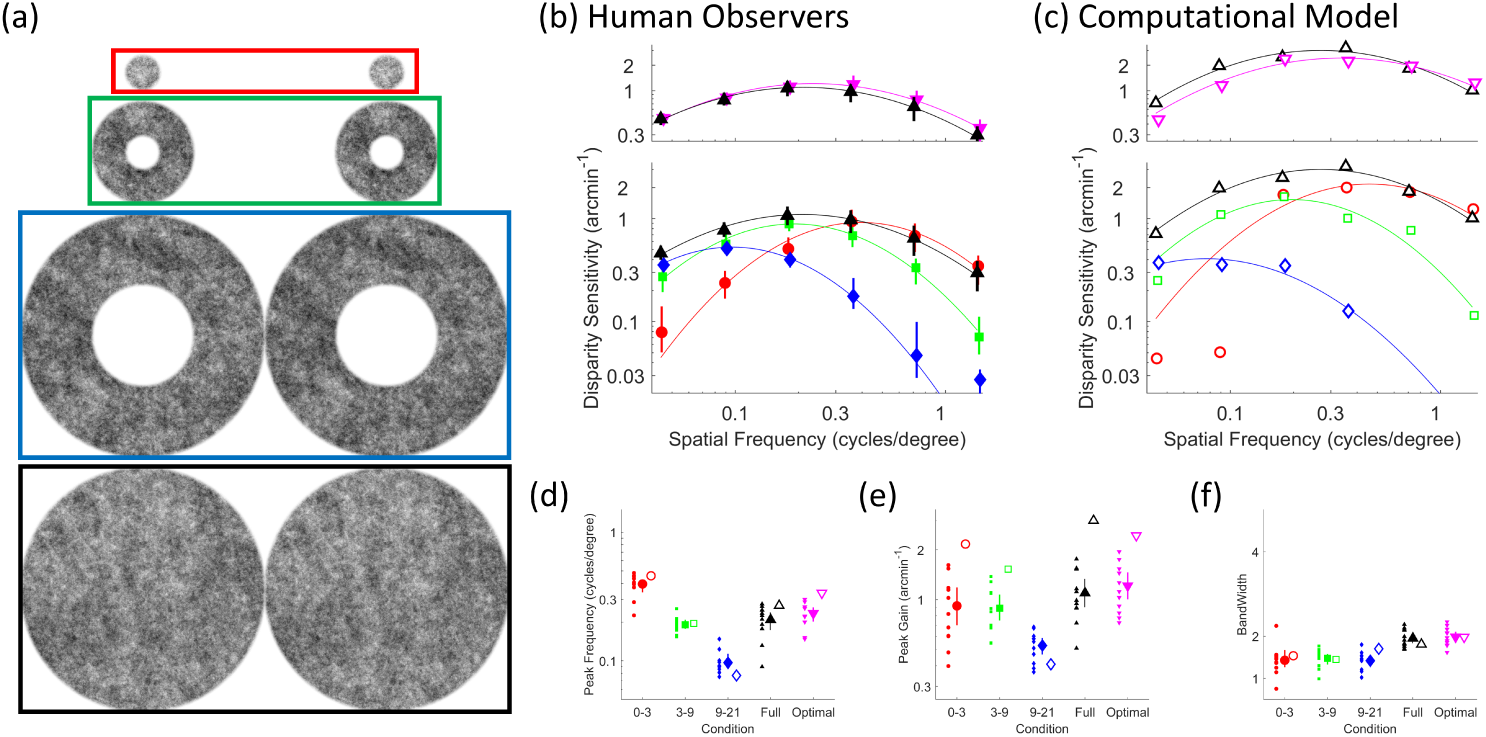
Disparity sensitivity across the visual field. (a) Participants and computational model were tested with annular pink noise stimuli spanning the foveal (red; 0-3 deg), mid (green; 3-9 deg), far (blue; 9-21 deg), and full (black, 0-21) visual field. (b) In the bottom panel, human disparity sensitivity is plotted as a function of spatial frequency for stimuli spanning far (blue diamonds), mid (green squares), foveal (red circles), and full (black upwards pointing triangles) portions of the visual field. In the top panel, human disparity sensitivity for the full field stimuli is compared to MLE-optimal disparity sensitivities (magenta downwards pointing triangles). Continuous lines are best fitting log parabola functions passing through the data (c) As in b, except for the computational model of disparity processing. (c-e) Peak frequency, peak gain, and bandwidth of the fitted log parabola model as a function of the portion of visual field tested, and for the MLE optimal sensitivity. In all panels, filled markers represent human data, empty markers represent data from the computational model of disparity processing. Small markers are data from individual participants, large markers are the mean sensitivities across participants and error bars represent 95% bootstrapped confidence intervals.

### Different regions of the visual field process disparity at different scales

Figure 1b (bottom plot) shows the tuning of human disparity sensitivity across different regions of the visual field. Disparity sensitivity in the far periphery (blue curve) is tuned to depth variations at low spatial frequencies. Disparity sensitivity in the near periphery (green curve) is tuned to depth variations at mid spatial frequencies. Disparity sensitivity in the fovea (red curve) is tuned to depth variations at high spatial frequencies. Thus, the peak frequency of the disparity sensitivity curves shifts from high to low frequencies moving from the fovea to the peripheral visual field (Figure 1d, *F*_2,18_ = 186.65, *p* = 9.2 × 10^-13^). The spatial frequency of depth variation at which peak sensitivity occurs also decreases from the fovea to the peripheral visual field (Figure 1e, *F*_2,18_ = 15.87, *p* = 1.1 *×* 10^-4^), whereas the bandwidth of disparity tuning remains constant (Figure 1f, *F*_2,18_ = 0.2, *p* = 0.82).

### Humans integrate disparity information across the visual field in a near-optimal fashion

Figure 1a (bottom plot) shows how disparity sensitivity for the full field stimuli (black) is the envelope of the disparity sensitivities estimated in the restricted visual field conditions. Additionally, Figure 1b (top plot) shows how disparity sensitivity for stimuli spanning the whole visual field (black) approaches the level of sensitivity predicted from the MLE optimal combination of disparity sensitivity across the separate portions of the visual field (magenta, following [22], see methods section for precise mathematical formulation). While qualitatively similar, disparity tuning for the full field stimuli was statistically different from the MLE optimal disparity tuning based on optmal integration of disparity across the retina. More specifically, disparity tuning for the full field stimuli exhibited lower peak frequency (Figure 1d, *t*(9) = 3.95, *p* = 0.0033) and lower peak gain (Figure 1e, *t*(9) = 2.67, *p* = 0.026) compared to the MLE optimal disparity tuning, but with a bandwidth that was not significantly different (Figure 1f, *t*(9) = 0.53, *p* = 0.61). Nevertheless, these differences amounted to a sub-optimal reduction in sensitivity of only 0.1 arcseconds, and a shift in tuning of only 0.02 cycles/degree.

### A foveated model of disparity processing accounts for the patterns of human data

Figure 1c shows the spatial frequency tuning of disparity sensitivity in our log-polar computational model of disparity processing, tested with the same stimuli and procedure as the human observers. This pattern is strikingly similar to the patterns of disparity sensitivity across the visual field of human observers (Figure 1b), and the model shows a high level of agreement with the human data (*r* = 0.91; *p* = 8.3 × 10^-10^; *r*^2^ = 0.83). Across experimental conditions, the estimates of peak frequency, peak gain and bandwidth for the computational model follow the same patterns as those of the human subjects, and cover a similar range (compare filled and empty symbols in Figure 1 panels d-e). The log-polar stage of the computational model is crucial for replicating the patterns of human data. A computational model without the log-polar processing stage exhibits markedly different patterns of disparity sensitivity and very low agreement with the human data (Supporting Figure S1; *r* = 0.046; *p* = 0.83; *r*^2^ = 0.0022). The non-log-polar model also shows how stimulus configuration cannot account for the observed patterns of human data. Contrary to what occurs in humans, in the non-log-polar model performance is best in the far peripheral condition where the model can integrate disparity information across the largest image area.

## Discussion

Our human behavioural data demonstrate that different regions of the visual field preferentially process disparity at different spatial scales. Our data broadly align with the shifts in spatial frequency tuning for depth reported by Prince and Rogers [14]. Furthermore, by approximately log scaling our stimuli, we show that the loss in peripheral sensitivity is not as steep as that found with equally-sized annular stimuli that, unlike our stimuli, do not compensate for the change in sampling density across the visual field. Therefore, contrary to the common intuition that depth processing is best at the fovea, our results show that disparity sensitivity depends on both spatial frequency and eccentricity. Disparity sensitivity to low and mid spatial frequencies is higher in the far and near periphery respectively, than in the fovea.

This change in tuning across the visual field, is remarkably similar to the change in naturally-occurring disparity statistics that have been reported for observers in natural indoor and outdoor environments [23]. We therefore speculate that the origin of this tuning may be related to the patterns of depth information the visual system has developed to process. We further observe that disparity information is integrated across the visual field in a near-optimal MLE fashion [16-22]. This finding informs how depth information at multiple scales is computed and combined across the visual field.

Of course, in the natural environment, the perception of depth does not rely exclusively on binocular disparity, but is supported by several sources of visual information, such as linear perspective and motion parallax, that are combined into a unified depth percept [17]. These different cues likely have different reliability across different regions of the visual field. For example, defocus blur is a more variable cue to depth than disparity near the fovea [29], but disparity is more variable than blur away from fixation [30]. Here, we have only shown that within a single cue, binocular disparity, depth information is integrated near-optimally across different regions of the visual field. It remains to be seen whether depth information within and among different sources, such as blur, perspective and disparity, can be successfully or optimally integrated across the human visual field. It also remains unknown whether such integration would be weighted by the different patterns of reliability for different depth cues. Nevertheless, the possibility that multiple cues are integrated is supported by the observation that experiencing congruent blur and disparity information across the visual field facilitates binocular fusion compared with incongruent pairings [31].

The pattern of human disparity sensitivity that we observe is well captured by our biologically-motivated model of disparity processing that critically incorporates the log-polar retino-cortical transformation. It is generally accepted that our visual system processes disparity along at least two [2, 4-10] or more [11-13] channels that are selective for depth changes at different spatial scales. A key insight provided by our work is that depth-selective channels emerge directly from the log-polar, retino-cortical transform, since log-polar spatial sampling acts as an “horizontal” multi-scale analysis, i.e by design it processes different spatial scales at different image locations [27, 32].

## Materials and methods

All methods were approved by the Internal Review Board of Northeastern University and adhered to the tenets of the Declaration of Helsinki.

### Disparity sensitivity in human observers

#### Participants

Author GM and nine naïve observers, (6 female, mean ± sd age: 24 ± 6) participated in the study. All subjects had normal or corrected to normal vision and normal stereo vision. Prior to testing, participants were screened using the Titmus stereopsis test and only participants with stereoacuity of 40 arcseconds or better were included in the study. Informed consent was obtained from all subjects.

#### Apparatus

The experiment was programmed with the Psychophysics Toolbox Version 3 [33, 34] in Matlab (MathWorks). Stimuli were presented on an BenQ XL2720Z LCD monitor with a resolution of 1920 × 1080 pixels (display dot pitch 0.311 mm) at 120 Hz. The monitor was run from an NVidia Quadro K 420 graphics processing unit. Observers were seated in a dimly lit room, 45 cm in front of the monitor with their heads stabilized in a chin and forehead rest and wore active stereoscopic shutter-glasses (NVIDIA 3DVision) to control dichoptic stimulus presentation. The cross talk of the dichoptic system was 1% measured with a Spectrascan 6500 photometer.

#### Stimuli

Stimuli were 1/f pink noise stereograms presented on a uniformly gray background; examples for each experimental condition are shown in Figure 1a. The stimuli contained oblique (45 or 135 degrees) sinusoidal disparity corrugations of varying amplitude and spatial frequency, generated as in [35]. The stimuli were presented as disks or rings with 1 degree cosinusoidal edges. The central fixation target was a 0.25 degree black disk with 0.125 degree cosinusoidal edge. In pilot testing, we verified that it was not possible to perform the experiment without dichoptic stimulus presentation (i.e. the oblique sinusoidal corrugation did not generate visible compression and expansion artifacts in the pink noise patterns).

#### Procedure

Each trial, observers were presented with a black fixation dot on a uniformly gray background. As soon as the response from the previous trial had been recorded, the stimulus for the current trial was shown for 0.25 seconds. Once the stimulus had been extinguished, observers were required to indicate, via button press, whether the disparity corrugation was top-tilted leftwards or rightwards. Observers were given unlimited time to respond. The following trial commenced as soon as observers provided a response. Each trial, the amount of peak-to-through disparity was under the control of a three-down, one-up staircase [36] that adjusted the disparity magnitude to a level that produced 79% correct responses.

#### Design

We measured how observer’s disparity sensitivity (1/disparity threshold) varied, as a function of the spatial frequency of the sinusoidal disparity corrugation, throughout different portions of the visual field. We tested four visual field conditions. In the central visual field condition, stimuli were presented within a disk with a 3 degree radius centered at fixation. In the near and far peripheral visual field conditions, stimuli were presented within rings spanning 3-9 and 9-21 degrees into the visual periphery, respectively. Lastly, in the full visual field condition, stimuli were presented within a disk with a 21 degree radius, and thus spanned the full extent of the visual field tested in this study. In each condition, we measured disparity thresholds at six spatial frequencies: 0.09, 0.18, 0.35, 0.71, 1.41, 2.83 cycles/degree. Thresholds were measured via 24 randomly interleaved staircases [36]. The raw data from 75 trials from each staircase were combined and fitted with a cumulative normal function by weighted least-squares regression (in which the data are weighted by their binomial standard deviation). Disparity discrimination thresholds were estimated from the 75% correct point of the psychometric function.

It is well known that disparity sensitivity varies lawfully as a function of spatial frequency following a bell-shaped function [37, 38]. This function is well described by a log-parabola model [35]. Therefore, we first converted disparity threshold estimates into disparity sensitivity (sensitivity= 1/threshold). Then, for each visual field condition, we fit the sensitivity data to a three-parameter log parabola Disparity Sensitivity Function (DSF) [35, 39] defined as:

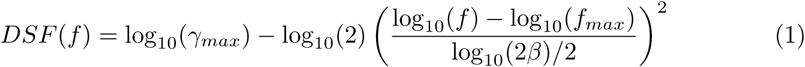

where *γ*_*max*_ represents the peak gain (i.e. peak sensitivity), *f*_*max*_ is the peak frequency (i.e. the spatial frequency at which the peak gain occurs), and *β* is the bandwidth of the function. The sensitivity data were fit to this equation, via least-squares regression, to obtain parameter estimates that could then be compared across experimental conditions.

### Ideal observer model

It is unknown whether observers are able to combine binocular disparity information across different portions of the visual field. If this were the case, then the DSF estimated for the full visual field condition should be the envelope of the DSFs estimated in the restricted visual field conditions. We obtained an estimate of the upper bound of performance in the full visual field condition by designing an ideal observer that optimally combines disparity information across the different portions of the visual field following a maximum-likelihood estimate (MLE) rule [22]. According to the MLE method, the disparity thresholds in the full-field condition should be lower (i.e. sensitivity should be higher) than in the the restricted visual field conditions, following the rule:

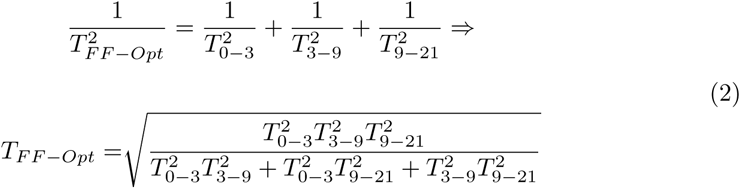

Therefore, we estimated the optimal disparity sensitivities as 1*/T*_*F* *F-Opt*_ at each tested spatial frequency. Then, we fit these optimal sensitivity data to the same DSF from Eq. 1 to obtain DSF parameter estimates for an ideal observer that could be compared to the DSF parameter estimates for the full field stimuli.

### Statistical Analyses

To test whether disparity sensitivity varied across the visual field of human observers, DSF parameter estimates from the restricted visual field conditions were analyzed using a one-way, within-subject Analysis of Variance (ANOVA). ANOVA normality assumptions were verified with Quantile-Quantile plots. Paired t-tests on the DSF parameter estimates were employed to test whether full field DSFs differed from MLE-optimal DSFs. To compare the computational model (described below) to human performance, we correlated the average human disparity sensitivity estimates to the model disparity sensitivity via simple linear regression.

### Foveated, image-computable model of disparity processing

We developed a biologically-inspired computational model that implements plausible neural processing stages underlying disparity computation in humans. The computational model mimics the dorsal visual pathway from the retinae to the middle temporal (MT) visual area [40, 41]. Critically, the model incorporates a biologically-plausible front end that approximates the space-variant sampling of the human retina. We hypotheses that this space-variant retinal sampling is responsible for the observed shifts in disparity tuning occurring across the visual field of human participants.

The computational model can be summarized as follows:

- a space variant front-end, i.e. the log-polar mapping that samples standard Cartesian stereo images;
- hierarchical neural processing layers for disparity estimation, based on V1 binocular energy complex cells and an MT distributed representation of disparity;
- a layer to take into account the optimal combination of disparity across annular regions of the visual field;
- a decoding layer in order to assess the encoded disparity into the cortical distributed representation.

Since the first processing stage is intended to mimic human retinal sampling, it consists of a log-polar transformation [24, 27] that maps standard Cartesian images onto a cortical image representation.

For disparity estimation we employ a feed-forward neural model that computes vector disparity [42]. This model can be directly applied on cortical images, since 2D vector disparity is computed without explicitly searching for image correspondences along epipolar lines. This allows us to discount the fact that straight lines in the Cartesian domain become curves in log-polar space [43], and this approach also does not require knowledge of the current pose of the stereo system (i.e. ocular vergence), even though in-principle this information could improve disparity estimation. It is also worth noting that even a simple (1D) disparity pattern is warped in the cortical domain. Therefore, to characterize properly a 1D Cartesian disparity pattern in cortical coordinates, a vector representation of cortical disparity is required.

To mimic the near-optimal combination of disparity information across different portions of the visual field of human participants, we consider a simple pooling mechanism that combines neural activity across annular regions of the model’s visual field.

To compare the model to human disparity processing, we decode the model’s distributed cortical activity and quantify the encoded disparity information. Even though this decoding stage is biologically plausible, we do not claim that it models the perceptual decision stage. We only employ this decoding stage to assess whether disparity estimation in the proposed model leads to patterns of disparity sensitivity similar to those measured in human participants.

### Retino-cortical mapping and cortical processing

To mimic the *retino-cortical mapping* of the primate visual system that provides a space-variant representation of the visual scene, we employ the central blind-spot model: each Cartesian image is transformed into its cortical representation through a log-polar transformation [24, 28, 44-46]. We chose this specific model with respect to other models in the literature (e.g. [47]) for several reasons: it captures the essential aspects of the retino-cortical mapping, it can be implemented efficiently, it provides a good preservation of image information [48, 49], and it allows us to provide an analytic description of cortical processing.

In the central blind-spot model, the mapping **T** : (*x, y*) ↦(*ξ, η*) from the Cartesian domain (*x, y*) to the cortical domain of coordinates (*ξ, η*) is described by the following equations:

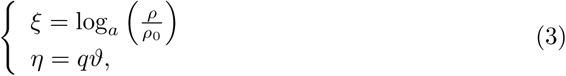

where *a* parameterizes the non-linearity of the mapping, *q* is related to the angular resolution, *ρ*_0_ is the radius of the central blind spot, and 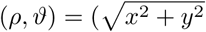, arctan (*y/x*)) are the polar coordinates derived from the Cartesian ones. All points with *ρ < ρ*_0_ are ignored (hence the central blind spot).

### Discrete log-polar mapping

Our aim was to test the model using the same experimental stimuli and procedures employed with human observers. Therefore, the log-polar transformation must be applied to digital images. Given a Cartesian image of *N*_*c*_ × *N*_*r*_ pixels, and defined *ρ*_*max*_ = 0.5 min(*N*_*c*_, *N*_*r*_), we obtain an *R* × *S* (rings × sectors) discrete cortical image of coordinates (*u, v*) by taking:

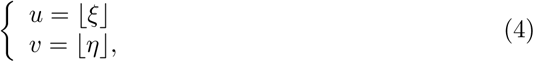

where ⌊ · ⌋denotes the integer part, *q* = *S/*(2*π*), and the non-linearity of the mapping is *a* = (*ρ*_*max*_*/ρ*_0_)^1*/R*^.

Figure 2 shows the log-polar pixels, which can be thought of as the log-polar receptive fields (RFs), in the Cartesian domain (b) and in the cortical domain (c): the Cartesian area (i.e. the log-polar pixel) that refers to a given cortical pixel defines the cortical pixel’s receptive field. The non-linearity of the log-polar transformation can be described as follows: by referring to Figure 2b and c, a uniform (green) row of cortical units is mapped to a (green) sector of space variant RFs, and a vertical (cyan) column of cortical units is mapped to a (cyan) circular set of uniform RFs. By inverting Eq. 3 the centers of the RFs can be computed, and these points present a non-uniform distribution throughout the retinal plane (see the yellow circles overlying the Cartesian images in Figure 2a). The magenta circular curve in Figure 2b, with radius *S/*2*π*, represents the locus where the size of log-polar pixels is equal to the size of Cartesian pixels. In particular, in the area inside the magenta circular curve (the *fovea*) a single Cartesian pixel contributes to many log-polar pixels (oversampling), whereas outside this region multiple Cartesian pixels will contribute to a single log-polar pixel. To avoid spatial aliasing due to the undersampling, we employ overlapping RFs. Specifically, we use overlapping circular Gaussian RFs [50, 51], which are the most biologically plausible and optimally preserve image information [48]. An example of a transformation from Cartesian to cortical domain is shown in Figure 2a and d. The cortical image (d) clearly demonstrates the non-linear effects of the log-polar mapping.

**Fig 2.**
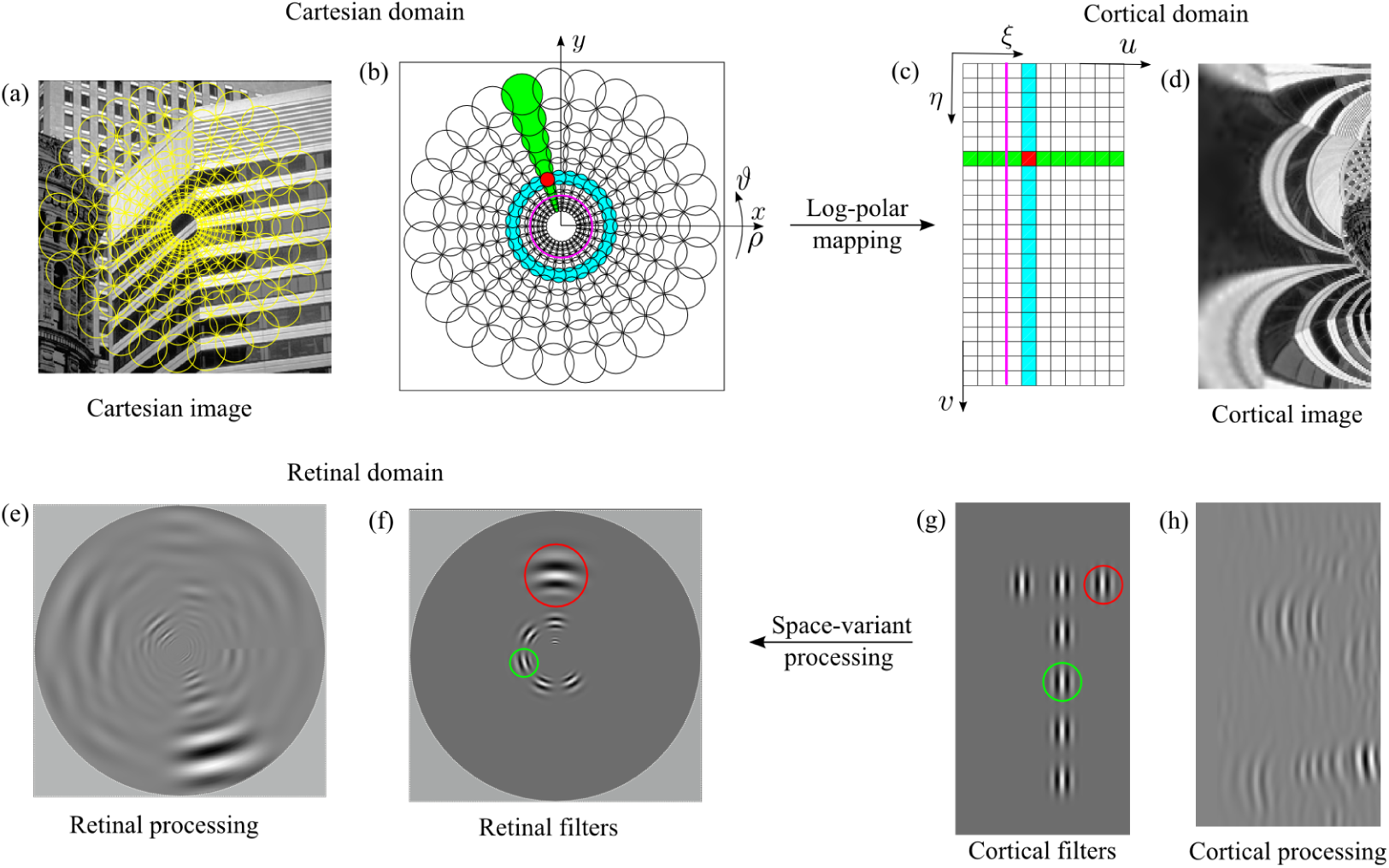
(top) Log-polar mapping scheme for the central blind-spot model (Eq. 3. (a) A standard Cartesian image with overlying log-polar pixels, the RFs (yellow circles). (b) Cartesian domain with the superposition of the circular overlapping log-polar receptive fields and (c) the corresponding cortical domain, where the squares denote the neural units. The green sector of RFs map to the horizontal row of (green) neural units and the cyan circle of RFs to a column of (cyan) neural units. The magenta circle delimits the oversampling (fovea) and undersampling areas (periphery). (d) The cortical representation of the standard Cartesian image. The cortical image is zoomed to improve the visualization. (bottom) A uniform processing in the cortical domain maps to a space-variant processing in the retinal domain. (a) The retinal space variant filtered image that is the backward mapping of the cortical uniform filtered image of subfigure (h). (f) The retinal filters (RFs) that correspond to the filters in the cortical domain (g): a uniform filtering in the cortical domain results in a space-variant filtering operation in the retinal domain, where both the scale (red circle) and the orientation (green circle) of the filters vary. (h) The cortical filtered image obtained by applying the filter depicted in subfigure (g) on the cortical image shown in subfigure (d). The specific values of the log-polar parameters are: *R* = 130, *S* = 203, *ρ*_*o*_ = 3, *CR* = 3.9, *W*_*max*_ = 4.8. The spatial support of the filter is 31 *×* 31 cortical pixels.

This discrete log-polar mapping provides a significant data reduction while preserving a large field of view and high resolution at the fovea [27, 52, 53]. To characterize the amount of data reduction provided by this transformation, we can can define the compression ratio (*CR*) of the cortical image with respect to the Cartesian one as:

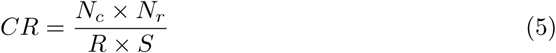

This compression ratio *CR* thus describes the data reduction occurring in the human visual system (that our computational model mimics), and will also affect the execution time of the simulated model.

The log-polar transformation models the space variant image resolution: the size of the RFs increases as a function of the eccentricity (the distance between the center of the RF and the fovea). We can define the relationship between the RF size (in particular, the maximum RF size *W*_*max*_) and the parameters of the mapping as follows:

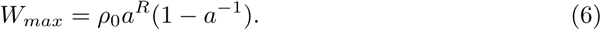

Equation 6 provides a measure of the scale at which the periphery of the Cartesian image is processed. Moreover, the parameters of the log-polar mapping also influence the proportion of cortical units used to over-represent the fovea: we can define the percentage of the cortical area used to represent the fovea (*χ*). This can be derived from Eq. 6 by setting the RF size to 1 and inverting the equation to find the corresponding *u* (see Eq. 4), and by then dividing by the overall size of the modeled cortex *R*:

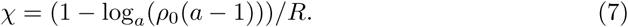

By exploiting Eqs. 6 and 7 we can control the growth of the size of the RFs and the over-representation of the fovea in order to reproduce data from the literature on the size-to-eccentricity relationship [46, 54, 55].

### Cortical processing

In the human visual system, visual processing is performed by networks of units (cells) described by their RFs. This neural network can be approximated by sets of filter banks whose responses to visual stimuli mimic those of neurons throughout the human visual system. The proposed model for disparity estimation could therefore embed the processing of V1 binocular simple units directly into the log-polar RFs. Specifically, the log-polar transform could be modified by using, as RFs, filters that perform V1-like feature extraction. However, to minimize the model’s computational load, we can consider that filter banks embedded in the log-polar transform can be “implemented” as a filtering process applied directly to the cortical image [27, 56]. We can demonstrate that the extraction of visual features can be carried out directly in the cortical domain by using solutions developed for the Cartesian domain without any modifications. To do so, in the following we analyze the relationships between the different parameters of a discrete log-polar mapping and of a bank of multi-scale and multi-orientation band-pass filters [57].

#### Sampling

To maintain equivalence between Cartesian and cortical visual processing, the discreet log polar mapping should provide an isotropic sampling of Cartesian coordinates. To avoid anisotropy, circular sampling must be (locally) equal to radial sampling, since the cortical space consists of a uniform network of neural units. Sampling points can be derived by considering the inverse of the cortical mapping **T** (Eq. 3). Specifically, the circular sampling interval is (2*π/S*)*ρ*_0_*a*^*u-*1^ and the radial sampling interval is *ρ*_0_*a*^*u-*1^(*a -* 1). To maintain isotropic sampling these sampling intervals must be equal, therefore the relationship between rings and sectors of the log-polar mapping must follow the rule:

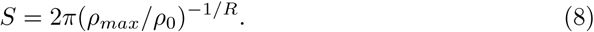

From a geometric point of view, the optimal relationship between *R* and *S*, expressed by Eq. 8, is the one that optimizes the log-polar pixel aspect ratio making it as close as possible to 1.

#### Receptive field shape

The RFs of V1 simple cells are classically modeled as band-pass filters [58], thus we define the following complex-valued Gabor filter [59]:

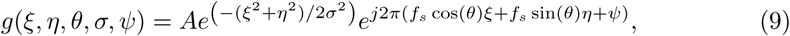

where *σ* defines the spatial scale, *f*_*s*_ the peak spatial frequency, and *ψ* is the phase of the sinusoidal modulation. By considering filters that are normalized by their energy, we have 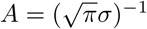.

In order to process the cortically-transformed images, it is necessary to characterize the filters, defined in the Cartesian domain, with respect to the cortical domain, i.e. to map the filters into the cortical domain, thus obtaining *g*(*x*(*ξ, η*), *y*(*ξ, η*), *θ, σ, ψ*). As a consequence of the non-linearity of the log-polar mapping, the mapped filters are distorted [60, 61], thus a filtering operation directly in the cortical domain could introduce undesired distortions in the outputs. Here, we show that under specific conditions these distortions can be kept to a minimum: under these assumptions, it is possible to directly work in the cortical domain, by considering spatial filters sampled in log-polar coordinates *g*(*ξ, η, θ, σ, ψ*),

At a global level (e.g. see Fig 2(d)) log-polar transformed images exhibit large distortions. However, we can consider what occurs at a more local level, at the scale of the RF of a single Gabor filter. First, we consider that the log-polar mapping can be expressed in terms of general coordinates transformation [62], thus the Jacobian matrix of the coordinates transformation allows us to describe how the RF locally changes. Specifically, the scalar coefficient *ρ*_0_*a*^*ξ*^ ln(*a*) represents the scale factor of the log-polar vector, and the matrix describes the rotation *η* due to the mapping. Figure 2g shows a set of cortical filters and Figure 2f their retinal counterpart (i.e. the inverse log-polar transform): the red circle highlights the scale factor (i.e. the spatial support) of the filter and the green one its rotation. It is worth to note that the column of “vertically” oriented filters in the cortical domain maps on a circle of filters in the retinal domain and each retinal filter is also at a different orientation.

Next, we want to analyze how the distortion affects the RF shape as a function of the distance from its center *p*_0_ = (*ξ*_0_, *η*_0_): we can consider that the ratio *g*(*x*(*ξ, η*), *y*(*ξ, η*), *θ, σ, ψ*)*/g*(*ξ, η, θ, σ, ψ*) around a given point should be equal to 1. Since the filter *g*(·) is an exponential function, we can evaluate the difference *h*(·) between their arguments. We can approximate such a difference by using a Taylor expansion of a multi-variable function:

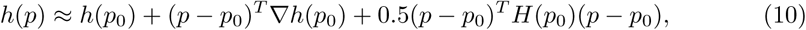

where (·)^*T*^ denotes the transpose, and *H*(·) the Hessian matrix. In the following we only focus on the terms that are relevant to describe how the distortion affects the RF shape: essentially, this depends on the partial derivatives of (*x*(*ξ, η*), *y*(*ξ, η*)) that constitute the gradient and the Hessian of *h*(·). The first order term takes into account how the mapping depends on the spatial position of the RF center. Indeed, the gradient has terms that are in common with the Jacobian matrix of the coordinates transformation, thus it describes the scale factor and the rotation of the RF as a function of the position *p*_0_. The approximation error can be expressed by the second order term of the Taylor expansion: thus, there is an error that increases as a quadratic function of the distance *p* − *p*_0_ (i.e. from the RF center), and an error that depends on the Hessian matrix that is related to the log-polar parameters. For instance, the mixed partial derivative of *x*(*ξ, η*) is *ρ*_0_ ln(*a*)*a*^*ξ*^ sin(*η*), thus we can consider that the error related to the log-polar parameters is proportional to *ρ*_0_ ln(*a*) = (*ρ*_0_*/R*) ln(*ρ*_*max*_*/ρ*_0_). It increases as a function of *ρ*_0_ (given a fixed *ρ*_*max*_) and decreases as *R* increases, which in turn decreases the compression ratio (Eq. 5). Figure 2f-g shows that such distortions can be negligible, though the spatial support of the displayed filters is large for sake of visualization. Figure 2h shows the cortical image (Fig. 2d) filtered by the filter that is drawn in different cortical positions in Figure 2g. In Figure 2e the retinal (i.e. space variant) processing is shown, which is obtained through the inverse log-polar mapping of Figure 2h.

#### Cortical computational model of disparity estimation

We consider a pair of (grayscale) cortical images *I*^*L*^(*p*) and *I*^*R*^(*p*), for all positions *p* = (*ξ, η*) that are the cortical representations of an input stereo pair of Cartesian images. Our goal is to define a computational model that is able to encode in its cortical activity the information related to the disparity present in the Cartesian images. The cortical images are a warped version of the Cartesian images. The representation of disparity is a vector quantity. We thus define the disparity map *δ*(*p*) = (*d*_*ξ*_, *d*_*η*_)(*p*) as the difference between the pair of cortical images at each position *p*. To compute this cortical disparity map, the proposed model is composed of several processing stages.

#### V1 Binocular energy computation and normalization

In the proposed model we consider two sub-populations of neurons at the V1 level: binocular simple cells and complex cells. V1 simple cells are characterized by a preferred spatial orientation *θ* and a preferred phase difference Δ*ψ* between the left- and right-eye components of a cell’s RF. We model the RFs of V1 simple cells as Gabor filters (see Eq. 9). The spatial support of the filters is defined as a function of their spatial radial peak frequency *f*_*s*_ and bandwidth 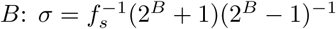. We consider one standard deviation of the amplitude spectrum as the cut-off frequency.

Following the *phase-shift model* [63, 64], we define the RFs of the binocular simple cell as *S*^*L*^(*p, θ, σ, ψ*^*L*^) = ℜ[*g*^*L*^(*p, θ, σ, ψ*^*L*^)] and *S*^*R*^(*p, θ, σ, ψ*^*R*^) = ℜ[*g*^*R*^(*p, θ, σ, ψ*^*R*^)]. These RFs are centered at the same position in the left- and right-eye images, and have a binocular phase difference Δ*ψ* = *ψ*^*L*^ − *ψ*^*R*^. For each spatial orientation, a set of *K* binocular phase differences are chosen to obtain tuning to different disparities: *d* = Δ*ψ/f*_*s*_.

We define the response of binocular simple cells as

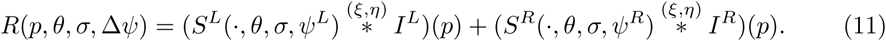

We can compute the response *R*_*q*_(*p, θ, σ,* Δ*ψ*) of a quadrature binocular simple cell by using the imaginary part of the Gabor filters.

The response of a complex cell is described by the binocular energy (the sum of the squared responses of a quadrature pair of binocular simple cells) [63, 65, 66]:

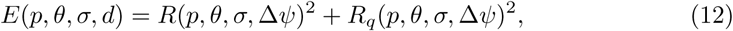

by considering that *d* = Δ*ψ/f*_*s*_. By taking into account the extensions of the binocular energy model proposed in [67, 68], we apply a static non-linearity to the complex cell response described in Eq. 12.

The response of the V1 layer of our model, when considering a finite set of orientations *θ* = *θ*_1_ … *θ*_*N*_, can be defined as

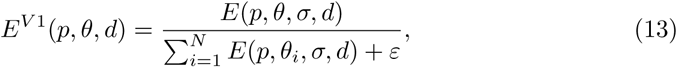

where 0 < *ε* ≪ 1 is a small constant to avoid dividing by zero in regions where no binocular energy is computed (i.e. no texture is present). For simplicity we omit from the notation the spatial scale *σ*. At this level, V1 responses are tuned to the spatial orientation and magnitude of the stimulus. In order to mimic natural neural activity, we consider that neural noise is present [67]. We model this neural noise as: *E*^*V*^ ^1^(*p, θ, d*) = *E*^*V*^ ^1^(*p, θ, d*) + *n*_*V* 1_(*p*). The noise is uniformly distributed and its value is a fraction of the local average neural activity.

#### MT cells response

Orientation-independent disparity tuning is obtained at the MT level of the model by pooling afferent V1 responses in the spatial and orientation domains, followed by a non-linearity [42, 69].

The responses of an MT cell, tuned to the magnitude *d* and direction *ϕ* of the vector disparity *δ*, can be expressed as follows:

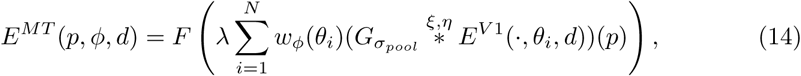

where 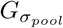 denotes a Gaussian kernel (standard deviation *σ*_*pool*_) for the spatial pooling, *F* (*s*) = *exp*(*s*) is a static non-linearity, specifically an exponential function [42, 70], *λ* is the gain of the non-linearity, and *w*_*ϕ*_ represents the MT linear weights that give origin to the MT tuning. Similarly to what occurs at the V1 layer, we model neural noise at the MT level as: *E*^*MT*^ (*p, ϕ, d*) = *E*^*MT*^ (*p, ϕ, d*) + *n*_*MT*_ (*p*).

Experimental evidence suggests that *w*_*ϕ*_ is a smooth function with central excitation and lateral inhibition. Therefore, by considering the MT linear weights shown in [70], we define *w*_*ϕ*_(*θ*) as

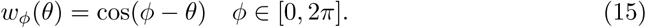

Vector disparity is thus encoded as a distributed representation through a population of MT neurons that span over the 2-D disparity space with a preferred set of tuning directions (*ϕ* = *ϕ*_1_ … *ϕ*_*P*_) in [0, 2*π*] and tuning magnitudes (*d* = *d*_1_ … *d*_*K*_).

Such a representation mimics the neural distributed representation of information. However, from a computational point of view, cosine functions shifted over various orientations (see Eq. 15) are described by the linear combination of an orthonormal basis (i.e., sine and cosine functions). Thus, all the V1 afferent information can be encoded by a population of MT neurons tuned to the directions *ϕ* = 0 and *ϕ* = *π/*2, only, with varying tuning magnitudes (see Eq. 14).

This description allows us to better understand how our model is able to account for a larger selectivity for the horizontal disparity, as reported in the literature [71-73]. Since a neural population tuned to two directions (at an angular difference of *ϕ* = *π/*2) can encode the full vector disparity, a neural population of MT units tuned in a range slightly larger than [− *π/*4, *π/*4] is able to recover the full vector disparity and shows also a larger selectivity for the horizontal disparity directions (i.e. cells tuned to directions around the horizontal one, in the retinal domain) [42].

#### Multi-scale analysis

A standard approach to handle multi-scale analysis is to adopt the following steps [42]: (i) a pyramidal decomposition with *L* levels [74] and (ii) a coarse-to-fine refinement [75]. This is a computationally efficient way to take into account the presence of different spatial frequency channels in the visual cortex and of large range of disparities and spatial frequencies in the real visual signal.

However, our model implements a log-polar mapping, thus its space variance, i.e. the linear increase of the filter size with respect to the eccentricity, can be exploited to efficiently implement a multi-scale analysis. Specifically, a pyramidal approach can be considered as a “vertical” multi-scale (the variation of the filter size at a single location), whereas the log-polar spatial sampling acts as an “horizontal” multi-scale (the variation of the filter size across different location [32]). The “vertical” multi-scale is also addressed in the literature as “cortical pyramids”.

#### Cue combination across the visual field

Human observers and model were tested with annular stimuli spanning sub-portions of the visual field, as well as with full field stimuli spanning the whole region of the visual field visual within a 21 degree radius. When analyzing the responses of the model to the foveal, mid-peripheral, and far-peripheral stimuli, we directly considered only the the activity (described by Eq. 14) of the neurons in the corresponding visual field regions. When analyzing the responses of the model to the full-filed stimuli, we instead pooled the neural activities of the distinct MT populations across the three considered annular regions.

#### Decoding

To assess whether the proposed computational model is able to effectively encode information about the features of the visual signal, and whether the model DSF is similar to the DSF of human observers, we decode the population responses of the MT neurons [67].

We adopt a linear combination approach to decode the MT population response as in [42, 76, 77]:

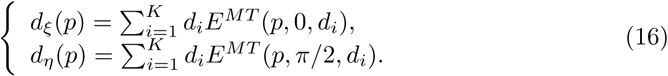

Then, we backwards transform into the retinal domain the disparity map described by Eq. 16. To easily detect whether the disparity corrugation is top-tilted leftwards or rightwards, we apply the Fourier transform to the retinal disparity map and check the position of the peak of its magnitude.

#### Simulation parameters

The simulation parameters selected to obtain the results presented in Figure 1 were adapted from the simulation parameters reported in [42], which were originally tuned to perform on computer vision benchmarks [78-81]. Since the proposed algorithm is meant to model human stereo vision, not compete on computer vision benchmarks, we modified the simulation parameters to reflect the known properties of the human visual system. The most notable differences are:

- The algorithm presented in [42] did not contain neural noise, which is instead present in the human visual system [28] and was thus incorporated into the current model
- In [42] a multi-scale approach was adopted with 11 sub-octave scales in order to recover a large range of disparities (common in computer vision) by using Gabor filters with peak frequency of 0.26 cycles/pixel. In the current model, only 2 scales were employed, since several authors have proposed two spatial frequency channels for disparity processing in humans [4-10]

The specific model parameters employed here were:

- *𝒟* = 1.52 pixels, the disparity range to which the neural units are sensitive (this range is constrained by the spatial peak frequency *f*_*s*_ of the filters)
- *K* = 5, the sampling of the disparity range, i.e. the number of neural units for a given spatial orientation *θ*
- the V1 static non-linearity is a power function with exponent 0.5
- *σ*_*pool*_ = 3.66 pixels, the spatial pooling of V1 responses (its standard deviation)
- *λ* = 0.65, the gain of the exponential static non-linearity at the MT level
- *N* = 12, the number of spatial orientations, i.e. the number of neural units that sample the spatial orientation *θ*
- the neural noise is set to 34% and 0.18% of the local average neural activity at the V1 and MT levels respectively
- *f*_*s*_ = 0.13 cycles/pixel, the radial peak frequency of the Gabor filters
- *σ* = 5.12 pixels, the standard deviation of the Gabor filters
- the Gabor filters are zero-mean
- *R* = 318, the number of rings of the log-polar mapping
- *ρ*_0_ = 9 pixels, the radius of the central blind spot
- *CR* = 6.4, the compression ratio of the cortical image compared to the Cartesian image

## Author Contributions

GM, MC, PJB and FS conceived and designed the study. GM and PJB collected the human participant data. GM analyzed the data. MC and FS developed the log-polar model of disparity processing. All authors wrote the manuscript.

## Acknowledgments

The authors thank Dr. Alexandre Reynaud for sharing experimental code employed to pilot this work. This research was supported by National Institutes of Health grant R01EY029713. Author G.M. was supported by a Marie-Sklodowska-Curie Actions Individual Fellowship (H2020-MSCA-IF-2017: ‘VisualGrasping’ Project ID: 793660).

## Data Availability Statement

Data and analysis scripts will be made available from the Zenodo database (doi:xx.xxxx/zenodo.xxxxxxx).

---

**Supporting Figure S1.**
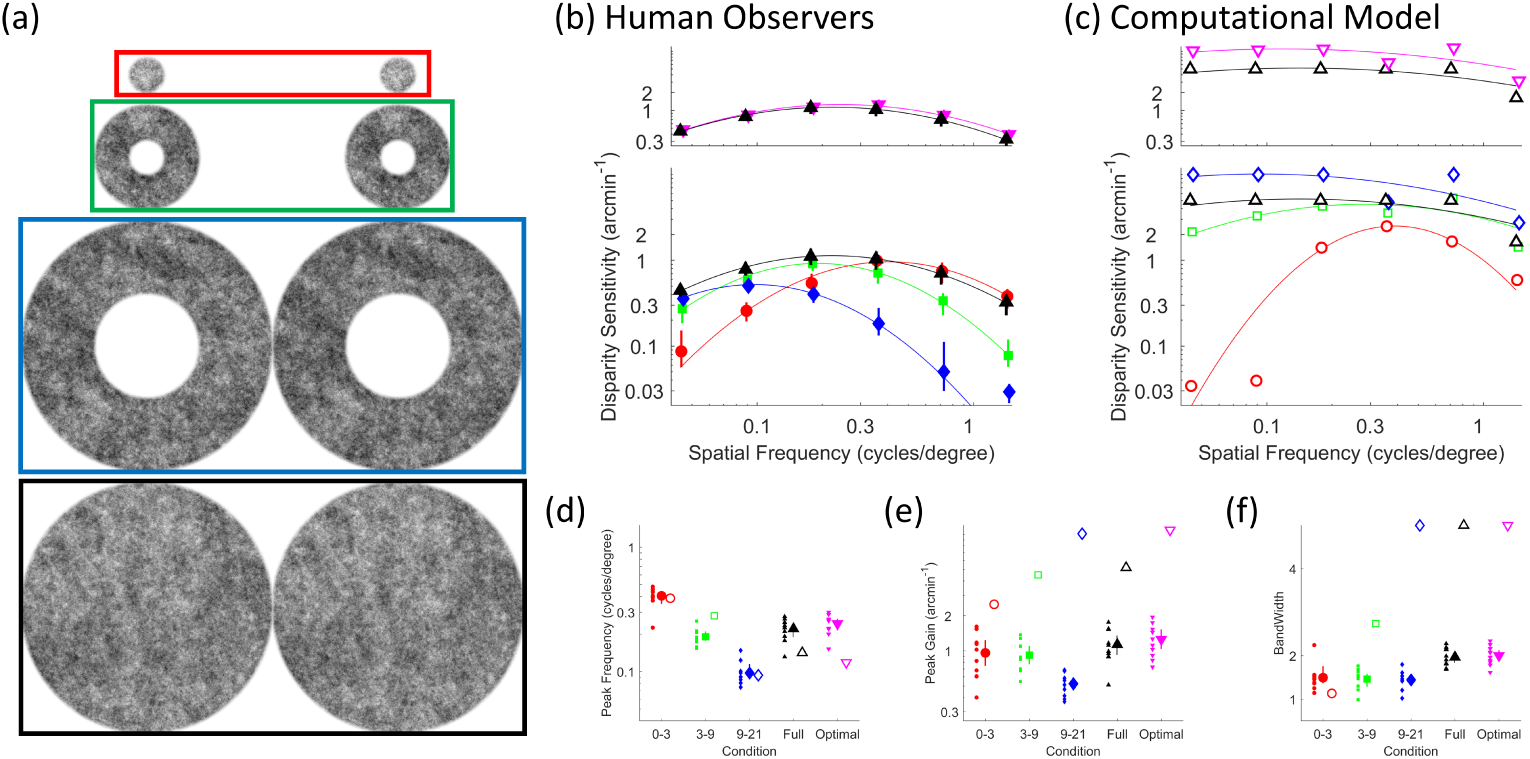
A computational model without the log-polar processing stage. Figure is constructed as Figure 1 from the main manuscript, except that the computational model does not contain a log-polar processing stage.

